# Vocal convergence during formation of social relationships in vampire bats

**DOI:** 10.1101/2025.06.21.660856

**Authors:** Julia Vrtilek, Grace Smith-Vidaurre, Eric Fosler-Lussier, Rachel A. Page, Gerald G. Carter

## Abstract

In many group-living birds and mammals, the formation of affiliative relationships is hypothesized to cause vocal convergence (an increase in call similarity between individuals). However, testing this causal effect can be difficult, because it requires experimentally forming new relationships. Here, we demonstrate convergence in the contact calls of common vampire bats (*Desmodus rotundus*) that we introduced and experimentally housed together. To estimate and disentangle the roles of kinship, co-housing (familiarity), allogrooming, and food sharing in predicting call similarity, we first measured call similarity using 35 features of 693,494 contact calls from 95 bats, then fit a series of Bayesian generalized multi-membership models. We also measured changes in call similarity for a subset of individuals that were recorded before and after co-housing. We found that co-housing caused vocal convergence. Furthermore, food-sharing rates among familiar nonkin within the same co-housed group predicted contact call similarity. This finding suggests that the development of cooperative relationships causes further vocal convergence beyond the initial convergence caused by co-housing. Our findings have implications for the development of cooperative relationships and vocal learning.

## Introduction

Vocal behaviour provides an important window into the social lives of many birds and mammals. Exchanges of vocalizations called “contact calls” seem to facilitate recognition, reunions, and collective movements among affiliated individuals across several species (e.g. African elephants (1), common marmosets (2), and white-winged vampire bats (3)). In many group-living birds and mammals, the contact calls of unrelated members of the same group sound more similar than those of strangers (e.g. black-capped chickadees (4), orange-fronted conures (5), budgerigars (6), yellow-naped amazons (7), pygmy and common marmosets (8–10), Guinea baboons (11), grey mouse lemurs (12), gelada (13), African elephants (14), bottlenose dolphins (15,16), orcas (17), greater spear-nosed bats (18) and Taiwanese leaf-nosed bats (19)). Based on this observation, a longstanding hypothesis is that the formation of affiliative relationships (“social bonds”) can increase similarity in the acoustic structure of calls between individuals (“vocal convergence”) (20).

Although vocal similarity or convergence is often associated with affiliative relationships (10,18,19,21–25), there are multiple possible mechanisms for how and why vocal convergence occurs. First, familiar animals might have the ability to modify the structure of vocalizations based on auditory inputs (vocal production learning), causing similarity between familiar individuals (e.g. 4). However, vocal convergence exists in many mammalian lineages where vocal production learning has not yet been demonstrated (26). In these cases, it is possible that animals might learn from conspecifics to produce existing sounds in their repertoire in new contexts (usage learning). Finally, familiar animals might use the same vocalizations more often simply because their physiological or motivational states are often matching (26).

Regardless of the underlying mechanism, demonstrating a causal link from affiliation to vocal convergence can be difficult. For example, correlations between affiliation and vocal similarity can result from kinship, either through heritable variation or shared environment. The same correlation can result from pre-existing vocal similarity causing higher affiliation (assortment based on acoustic similarity). To demonstrate causation, an ideal study would induce new relationships to form among nonkin, then observe changes in call similarity (e.g. 4).

Here, we applied this approach to demonstrate convergence in the contact calls of common vampire bats (*Desmodus rotundus*). Vampire bats produce contact calls when they are socially isolated (27) and use the calls to vocally recognize affiliated individuals (28). By analysing 693,494 contact calls from 95 bats, some of which were experimentally co-housed together, we found that co-housing causes vocal convergence. Next, to tease apart how pairwise call similarity was shaped by relationships within each captive group, we estimated effects of genetic relatedness, allogrooming, and food sharing (29–32). Both kin and nonkin females can develop reciprocal allogrooming relationships that can become reciprocal food-sharing relationships, in which a fed bat regurgitates potentially life-saving ingested blood to a starving partner (29–31). We found evidence that these food-sharing relationships cause further vocal convergence between nonkin after the initial convergence caused by co-housing.

## Methods

### Study subjects

The 95 common vampire bats (*Desmodus rotundus*) used in this study came from 8 different origins (e.g. captive colonies or field sites; for details, see Supplementary Information (SI), Table S1). To identify individual bats, all bats were marked using 1-4 unique bat bands. We labelled bats as “adults” if they were older than 2 years old at time of recording or, if their exact age was unknown, by the presence or absence of an epiphyseal gap between their metacarpals. We refer to volant non-adults as “subadult”. Non-volant pups that were still nursing from their mother were not included in the study.

### Kinship, social familiarity, and cooperative relationships

Bats captured from different sites hundreds of kilometres apart were considered both unrelated (non-kin) and unfamiliar. Kinship of bats from the same colony was estimated through a combination of microsatellite markers and maternal pedigrees, as described in previous studies (29,31). Pairs of bats with unknown kinship were excluded from analyses where kinship was a predictor.

Most pairs (n = 2605) were unfamiliar because they had never met; 582 pairs were captured from different field sites hundreds of km apart, then introduced in captivity and caged together (co-housed) for at least 4 months; 612 pairs were captured from the same field site then caged together for at least 4 months; and 435 pairs were living in captivity for years. Another 231 pairs had unknown levels of kinship and familiarity, because they were captured from the same wild roost about four years apart. Within each co-housed captive group, pairs had variable rates of cooperative behaviour. We had measures of food-sharing rates for 307 pairs, allogrooming rates for 190 pairs, and both measures for 406 pairs. These data were collected from a series of past studies where experimenters repeatedly induced food sharing by removing a bat, fasting it for over 24 hours, and reintroducing it to its home cage, then used focal follow sampling to observe allogrooming, regurgitated food sharing, or both (29–31).

### Recording contact calls from individually isolated bats

Contact calls are frequency modulated syllables that are produced by socially isolated bats (27). To record contact calls, we isolated an individual bat inside a plastic storage bin (about 75 to 200 litres in volume) lined with acoustic foam and/or soft fabric to dampen echoes. To keep the bat within 10 to 30 cm of the microphone, we placed it inside a small soft-mesh butterfly cage or a tube of plastic mesh. Calls were recorded with an Avisoft CM16 ultrasound condenser microphone (frequency range 10-200 kHz, purchased prior to 2017 from Avisoft Bioacoustics, Berlin, Germany). Sound was digitised with 16-bit resolution at sampling rates ranging between 250-500 kHz through an Avisoft UltraSoundGate (one-channel or four-channel) to a laptop running the program Avisoft Recorder. To remove any effects of digitisation rate on subsequent call measures, recordings digitised at a rate above 250 kHz were downsampled to 250 kHz using the *seewave* R package (33). To filter out sounds that were not contact calls, we removed calls with durations less than 3 ms or longer than 50 ms and with peak frequencies below 10 kHz. For further explanation of call selection, see SI, Supplementary Methods.

We analysed 693,494 contact calls from 95 of the 115 common vampire bats that were individually isolated and recorded from 2011 to 2019. We only analysed the 95 bats that produced at least 100 contact calls of sufficient quality to measure acoustic features (described below). The number of recording sessions per bat varied from 1 to 22 (mean = 7.36). The mean number of calls recorded per bat was 7300 (median = 2500, minimum = 118, maximum = 98161).

### Measuring acoustic features of calls

We wrote a custom R script to identify start and end times of individual calls using both amplitude and spectral density (the distribution of amplitude across frequencies). This automated method was tested and validated with 1535 manually labelled recordings containing ∼7000 call selections. Using these start and end times, we then measured 35 features of each call (Figure 1, Table S2). We used the R package *warbleR* (34) to measure 27 spectral and temporal parameters and the R package *soundgen* (35) to estimate 8 additional measurements from the fundamental: maximum slope, minimum slope, and absolute minimum slope; the number of positive slopes and of negative slopes; the number of turns; the mean slope; and the number of segments. We eliminated ‘clipped’ calls, which we identified as calls that reached a relative amplitude of more than 0.99 using the *normalize* function from the R package *tuneR* (36). All scripts were run using the Ohio Supercomputer (37).

**Figure 1.**
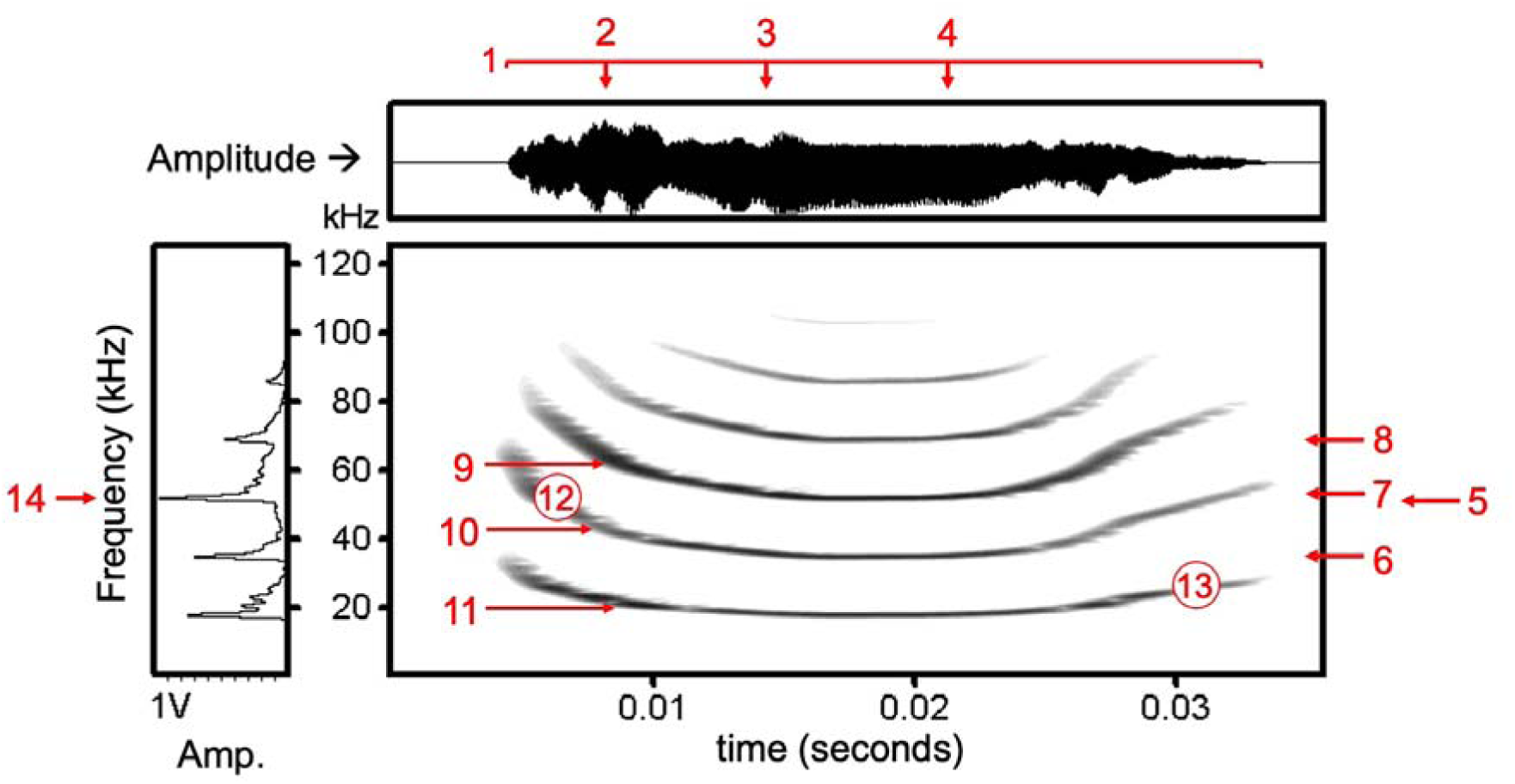
Vampire bat contact call. Call measures shown here include: 1 duration, 2 time quantile 25, 3 time median, 4 time quantile 75, 5 mean frequency, 6 frequency quantile 25, 7 frequency median, 8 frequency quantile 75, 9 maximum dominant frequency, 10 mean dominant frequency, 11 minimum dominant frequency, 12 start dominant frequency, 13 end dominant frequency, 14 peak frequency. Table S2 defines the above measures and 21 additional measures not shown here.

### Assessing vocal individuality

To assess and confirm baseline individuality in contact call structure, we estimated the classification rate for assigning calls to the correct caller using a linear discriminant function analysis (DFA) with leave-one-out cross-validation, including all variables measured (via the MASS R package (38)). To choose a minimum number of calls per bat, we compared these classification rates to the same rates from a single DFA without cross-validation. We excluded bats with fewer than 100 sampled calls because within-bat classification rates were virtually identical for bats with greater than 100 calls. To confirm that calls could be assigned to the caller with greater-than-chance accuracy, we ran a permutation test: we repeated a DFA without cross-validation 1000 times with randomized bat names to generate an expected distribution of accuracy rates under the null hypothesis that calls contain no information about caller identity.

### Estimating contact call similarity

We defined contact call similarity for each pair of bats as:

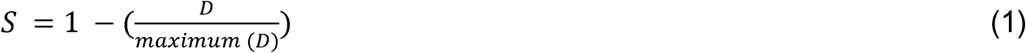

where D is the Mahalanobis distance between the two group centroids (bats) from a DFA classifying calls (n = 693,494) to bat (n = 95). These Mahalanobis distances represent the average multivariate distance (or dissimilarity) between every two bats. We normalized our measure of call similarity to range from 0 to 1; however, we replaced the single similarity score of 0 with 0.001 (the next lowest call similarity was 0.03), because similarity cannot be precisely zero and beta regression cannot model an outcome of zero.

### Estimating effects of kinship and social relationship on call similarity

To assess predictors of contact call similarity while controlling for the effect of both bat identities, we fit five Bayesian generalized (beta distribution) multi-membership models (Table 1) using the R package *brms* (39). Multi-membership models are used to model dyadic data by fitting group-structured random effects, which capture the dependence of each pair on the identities of both individuals. This model structure allows us to estimate a random intercept for each bat and its influence on its pairings with other bats (40). To assess causal hypotheses, we estimated multiple predictors across seven models based on known and hypothesized causal paths (Figure 2a).

Our first goal was to disentangle the effects of kinship from social familiarity. In model 1, we estimated how well kinship predicted contact call similarity across the 2179 pairs of adult bats with known kinship estimates. Kinship is confounded with group-level familiarity and within-group affiliation rates, and any effects of kinship in model 1 could therefore be driven by social interaction history rather than kinship. In model 2, we therefore estimated how well kinship predicted contact call similarity when conditioning on (i.e. statistically controlling for) affiliation rates in the same 2179 pairs. Predictors were kinship, whether the bats were co-housed (true/false), and within-group affiliation lograte, defined as:

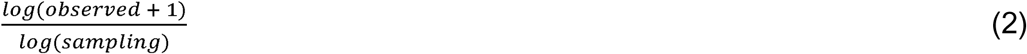

where “observed” was the seconds of either allogrooming or regurgitated food sharing between the two bats and “sampling” was the total seconds in which allogrooming or food sharing could have been observed between the two bats. Both allogrooming and food sharing are directed behaviours that indicate an affiliative social preference. We used logrates rather than rates because seconds of allogrooming and food sharing are lognormal. We scaled (i.e. z-transformed) both kinship and affiliation lograte to convert units to standard deviations.

Another way to control for kinship is to only test unrelated pairs. In model 3, we measured how well being caged together (true/false) predicted contact call similarity in the 1159 pairs of unrelated female adults caught from different sites. All pairs had kinship estimates of 0, allowing us to test for an effect of nonkin being caged together on their contact call similarity. We focused on female pairs because the few males in these colonies were sub-adults or young born into the group.

Our next goal was to separate the effects of group-level familiarity and dyad-level affiliation rate among nonkin females. In model 4, we measured how well within-group affiliation lograte (scaled lograte) predicted contact call similarity in 272 unrelated pairs of female adults that were caged together. We removed pairs with kinship estimates greater than 0.05 to test for an effect of nonkin social interactions on contact call similarity. We also removed pairs that could never have been in contact. We again focused on female pairs because social affiliation data were focused on females and undersampled or missing for males (29–32). We did not include allogrooming and allofeeding together as covariates in model 4 because they are highly correlated proxies for social affiliation, rather than different effects. The affiliation rate therefore includes either allogrooming, regurgitated food sharing, or both. The results of model 4 must therefore be interpreted carefully, because affiliation rate is based more on allogrooming in some pairs than others. For example, in one colony from 2010-2014, food-sharing rates were sampled more than allogrooming, but in another colony in 2019, food-sharing rates were unknown (missing) for all pairs. Moreover, food sharing represents a much higher level of affiliation than allogrooming. For example, when bats are introduced in captivity (i.e. become familiar), most pairs form allogrooming relationships and only a few begin food-sharing relationships (29). In other words, co-housing predicts allogrooming better than food sharing.

To remedy these issues, we used our last model to focus on food-sharing rates, which is the best indication of a within-group social bond (29). In model 5, we measured how well food-sharing rate (scaled lograte) predicted contact call similarity in the 139 unrelated pairs of female adults that were caged together. We again removed pairs with kinship estimates greater than 0.05, pairs that could never have shared food and pairs with unknown food-sharing rates.

Across all models, we used default priors and scaled continuous predictors. We used posterior predictive checks to assess model fit (Figure S2). To ensure Markov Chain Monte Carlo (MCMC) convergence, we checked the R-hat values across 4 chains, which ranged from 0.9999 to 1.0004, showing that all 4 chains reached the same estimates for coefficients. We used chain and warmup lengths of 6000 and 2000 samples, respectively. The lowest Bulk and Tail effective sample sizes across all models were 8747 and 10064 respectively.

**Table 1.** Purpose of each statistical model. We selected this set of models based on causal models shown in Figure 2a.

| Model | Question addressed | Covariates included as predictors of contact call similarity | Bat pairs in sample |
| --- | --- | --- | --- |
| 1 | How well does kinship predict contact call similarity? | kinship | pairs with estimated kinship = 2179 |
| 2 | How well does kinship predict contact call similarity after controlling for co-housing and affiliation? | kinship, co-housing (true/false), affiliation rate (including zeros for bats that never met) | pairs with estimated kinship = 2179 |
| 3 | In nonkin females, how well does co-housing predict contact call similarity? | co-housing (true/false) | pairs of nonkin adult females = 1159 |
| 4 | In nonkin females, how well does affiliation predict contact call similarity? | within-group affiliation rate (excluding bats that never met) | pairs of nonkin adult females that were caged together = 272 |
| 5 | In nonkin females, how well does food sharing predict contact call similarity? | within-group food-sharing rate (excluding bats that never met) | pairs of nonkin adult females with estimated food-sharing rates = 139 |

### Comparing call similarity pre-introduction and post-introduction

We could determine how call similarity changed over time using a subset of cases in 2016 where 7 bats from Tolé were recorded both before and after being introduced to 5 bats from Las Pavas (Figure S3). We predicted that contact call convergence (increases in call similarity) should be greater for introduced pairs (from different sites) than for previously familiar pairs (from the same site). To compare call similarity before and after introduction, we first performed two discriminant function analyses (DFAs) classifying calls to bats. Both DFAs included 5 Las Pavas bats and 7 Tolé bats, but the first DFA included recordings from the Tolé bats *before introduction* to the Las Pavas bats, and the second DFA included the same Tolé bats recorded *after the introduction*. From each DFA, we extracted Mahalanobis distances and call similarities as described above. We then estimated the change in similarity as the difference between post-introduction and pre-introduction call similarities.

To test the hypothesis that increases in similarity were greater for introduced pairs than for familiar pairs, we used two approaches. First, we used a nonparametric permutation test (Mantel test with Spearman correlation) of the correlation between the change in contact call similarity and a binary familiarity matrix indicating familiar vs. introduced pairs. Second, we used a Bayesian multi-membership model (Gaussian distribution, default priors in brms, 4 chains with length of 6000 interactions, 2000 warmup samples) to test whether familiarity predicted the change in contact call similarity.

Finally, we tested the prediction that call classification to the correct capture site should be more accurate prior to the merging of the two groups. To do this, we used two permuted DFAs, which are designed to test for correct classification of calls to social group (41). In the first permuted DFA, we classified calls to either the Las Pavas group or the *pre-introduction* Tolé group. In the second permuted DFA, we classified calls to either the Las Pavas group or the *post-introduction* Tolé group.

## Results

### Contact call variation within and between individuals

Contact calls had clear individual signatures but were also highly variable within bat (Figure S1). Discriminant function analysis (DFA) correctly assigned the 693,494 calls to the 95 callers with 36% probability, which is more than twice the expected chance level of 14% (95% quantiles of expected values: 14.14% to 14.15%). The call assignment accuracy of assigning calls back to individual bats from the cross-validated DFA and from the non-cross-validated DFAs were almost perfectly correlated across the 95 bats (r = 0.9996), indicating robustness of the DFA used for estimating call similarity.

### Kinship-biased contact call similarity can be explained by social familiarity

Model 1 revealed that kinship predicts contact call similarity (Figure 2b, Figure S4). However, kinship is confounded with social affiliation in vampire bats (30,31,42). We therefore used Model 2 to test for an effect of kinship while conditioning on (i.e. “controlling for”) affiliation rate and co-housing. We no longer found an effect of kinship while conditioning on these measures of social affiliation, but we did find that call similarity was predicted by both affiliation rate and co-housing while conditioning on kinship and each other (Figure 2b). This result does not mean that kinship did not influence call similarity; only that the correlations between kinship and contact call similarity can also be explained by social familiarity. In other words, both affiliation rate and familiarity predicted contact call similarity, regardless of kinship.

### Co-housing causes contact call similarity among nonkin

To further test whether co-housing causes contact call convergence among nonkin, we used Model 3 to focus on 1159 pairs of unrelated female adults that were captured from different sites. Compared to model 2, which included kin, we found that co-housing was associated with an even larger increase in call similarity among pairs of unrelated adult females (Figure 2b). As expected, call similarity for nonkin from distant sites that were housed together in captivity was dramatically higher than nonkin from different sites that never met, and the highest mean similarity in nonkin contact calls occurred within nonkin pairs from the same wild roost or in the same long-term captive colony (Figure S5).

### Within co-housed groups, food-sharing rates predict further vocal convergence

Although Model 2 revealed an effect of affiliation (allogrooming and food sharing, Figure S6), further analyses showed that this effect was driven by food sharing rather than allogrooming (Figure 2b, Figure S7). Specifically, Model 4 estimated the effect of within-group dyadic affiliation rate on call similarity among unrelated female adults, and this model found no clear effect of affiliation rate beyond the effect of previous familiarity. However, Model 5 showed that contact call similarity was predicted by within-group rates of regurgitated food sharing. This effect of food sharing on call similarity was largely driven by the 2014 long-term captive colony (Figure S7), which had the highest rates of food sharing and the longest periods of familiarity.

**Figure 2.**
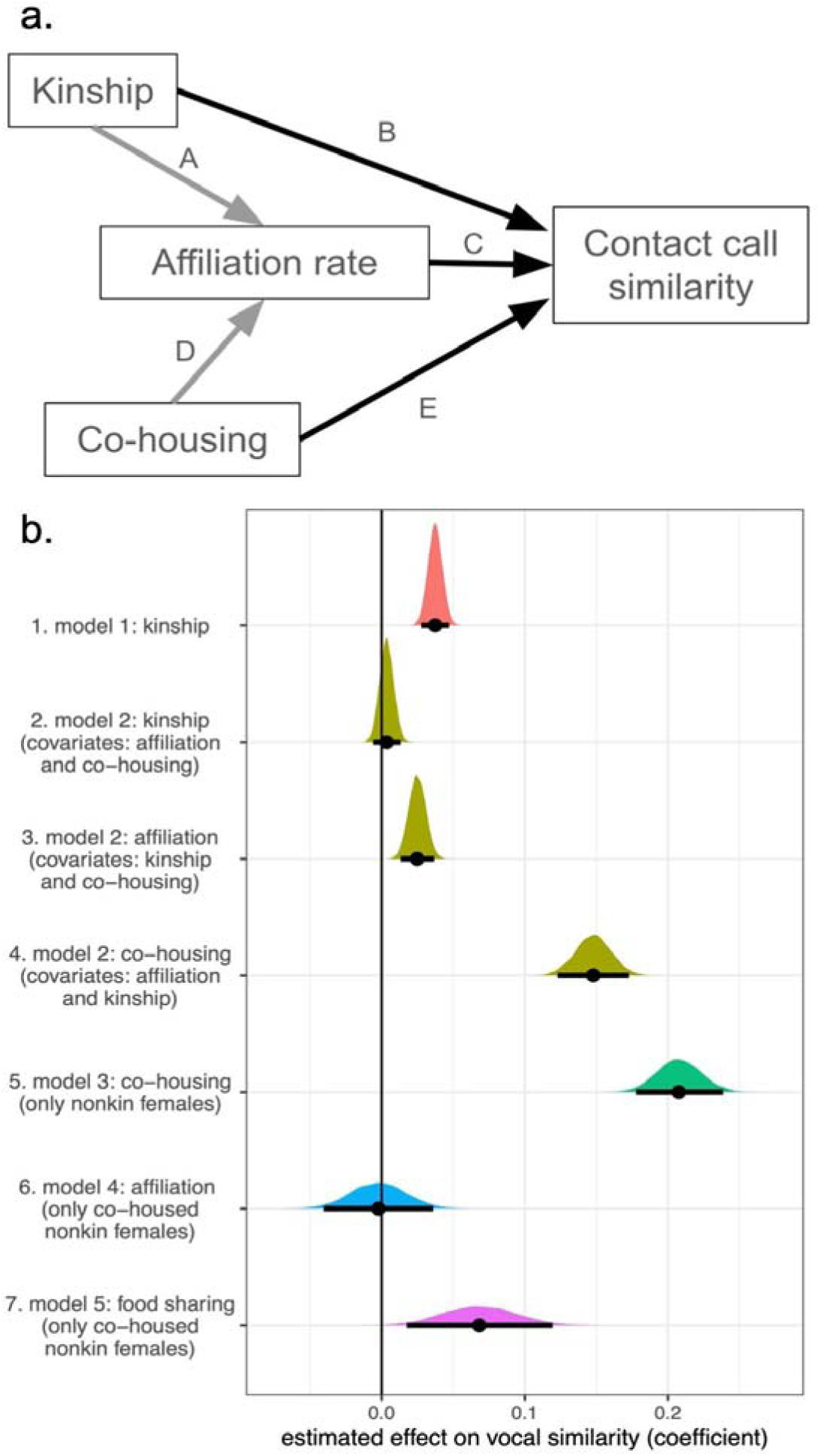
Contact call similarity in vampire bats is predicted by co-housing, affiliation rate, and food-sharing. **Panel a**: causal model for vocal convergence, with black arrows indicating causal paths of interest, and grey arrows indicating causal paths that cause potential confounding. Path A: Greater kinship causes greater affiliation. We estimate affiliation through allogrooming and food-sharing. Path B: Kinship might cause greater similarity in contact calls if call features are heritable. Path C: Affiliation might cause vocal convergence. Path D: Affiliation is caused by co-housing. Path E: Co-housing leads to acoustic exposure, which might also cause vocal convergence independent of variation in affiliation. **Panel b:** seven regression coefficients from five models (colours) selected to disentangle causal effects of kinship, co-housing, affiliation rate, and food-sharing. Dots and error bars show means and 95% credible intervals from Bayesian posterior probability distributions. See Table S3 for coefficients and credible intervals.

### Vocal convergence evident from call similarity before and after co-housing

We saw evidence for vocal convergence over time (increased contact call similarity after introduction) in 25 of 35 pairs from different sites (71%), compared to 11 of 21 familiar pairs from Tolé (52%). We found that a greater increase in call similarity (i.e. greater vocal convergence) was positively associated with being an introduced pair of bats from different sites (Mantel test: Spearman’s correlation = 0.28, n = 12 bats, p = 0.011; Bayesian multi-membership model: coefficient for being introduced = 0.06, Bayesian 95% credible interval = 0.02 - 0.11). Consistent with vocal convergence, the ability to correctly classify calls to capture site (Las Pavas or Tolé) decreased from 72% to 65% after bats from both capture sites were merged into one colony.

## Discussion

### Variation in social affiliation drives convergence of contact calls in vampire bats

For this study, we analysed a nine-year collection of contact calls recorded from a large sample of vampire bats for which we had known kinship and social interaction histories. We observed multiple cases of contact call convergence within both kin and non-kin pairs with different levels of familiarity (from completely unfamiliar to long-term co-housing) and relationship development (mere familiarity to reciprocal food-sharing relationship). Using these data, we demonstrated that familiarity, which was induced by co-housing bats in captivity, causes vocal convergence. We showed that the correlation between kinship and call similarity can be explained by social affiliation among kin.

Interestingly, food-sharing rates among familiar nonkin within the same co-housed group predicted contact call similarity, even beyond the effect of familiarity caused by co-housing. These findings are the first clear evidence of vocal convergence in vampire bats, and the first evidence of vocal convergence in bats based on within-group variation in cooperative relationships.

The mean levels of call similarity among nonkin were exactly what one would predict if familiarity causes vocal convergence (Figure S5). We observed the lowest contact call similarity among nonkin that never met and clearly higher call similarity among nonkin that were captured from different wild roosts and then co-housed in captivity. We observed the highest levels of call similarity equally among wild-caught nonkin from the same roost that were co-housed together, and among captive-born nonkin co-housed in the same long-term captive colony for years (Figure S5).

Food-sharing was associated with vocal convergence, but unexpectedly, we found that call similarity among co-housed bats was not predicted by allogrooming. This finding was clear from the difference between the effect of overall affiliative behaviour (allogrooming and food sharing) and the effect of only food-sharing rate (Figures S7 and S8). One interpretation of this result is that introduced (co-housed) bats vocally converge to some degree, and then additional convergence occurs for the subset of bat pairs with even stronger bonds (i.e. indicated by frequent food sharing). Compared to food-sharing relationships, allogrooming relationships are less stable and less selective.

All food-sharing relationships include allogrooming but not all allogrooming relationships involve food sharing, and when we co-housed previously unfamiliar adult female vampire bats for almost a year, 78% of those pairs began allogrooming while only 16% of them began food sharing (29). Moreover, the allogrooming relationships in this study were changing more than in a typical colony due to sickness behaviour from infections (43) and experimental manipulations such as forced proximity (29,32). Our analyses averaged these complex dynamics over the course of an entire study into a single mean pairwise allogrooming rate, which might not necessarily reflect the observed call similarity for the two bats that were each recorded at different times. In contrast, food-sharing rates in a long-term captive colony were reliable, strong, and stable indicators of a close social bond.

### Variation in affiliation and vocal convergence in other species

We observed vocal convergence in vampire bats at two levels: group-level familiarity (co-housing) and within-group close social bonds (food-sharing relationships). We believe this to be the first evidence of partner-specific vocal convergence in bats, in contrast with convergence on a group-level signature, which has been observed in the harsh broadband “screech” contact calls of greater spear-nosed bats (*Phyllostomus hastatus*) (18) and the tonal constant-frequency echolocation calls of Taiwanese leaf-nosed bats (*Hipposideros terasensis*) (19). Outside bats, partner-specific vocal convergence has been reported in multiple birds and mammals, including in bottlenose dolphins (15,44), orange-fronted conures (45), African elephants (46), and Campbell’s monkeys (23).

Vocal convergence at multiple levels of association (e.g. relationship and group) has been observed in several lineages of avian and mammalian vocal learners. In budgerigars, males entering a new flock adapt their social call to match the group (47), and mated pairs develop a shared call (48). Monk parakeets show acoustic convergence at the pair, flock, and site social scales (49). In crossbills, ecologically diverged but sympatric groups have discrete call variants, and some bonded pairs produce nearly identical call structures (50). In mammals, evidence of multi-level vocal convergence has come from cetaceans (e.g. 15,16,44) and nonhuman primates (e.g. 8,11,24). Bottlenose dolphins have individual signature whistles (15,44), but one group has also been shown to share a stereotyped whistle contour (16). Male Guinea baboons live in a multilevel society and individuals have more similar grunts within gangs (the largest group size) than between gangs, and within a party (several parties to a gang) than between parties (11). In the pygmy marmoset *Cebuella pygmaea*, there is strong evidence for convergence at the pair level and weak evidence at the group level: individuals converge the duration and structure of their trill calls in response to pairing with a new mate, but only converge duration when introduced to a group of strangers (8,24). In general, vocal convergence at multiple levels of social familiarity may be widespread, but there is not yet a clear pattern in how or why it is used by callers and receivers.

### Is vocal convergence an adaptive social behaviour or a byproduct of vocal plasticity?

Many authors argue that vocal learning requires complex and therefore costly neurological adaptations, and is therefore unlikely to evolve unless there is a significant fitness advantage (51). One possible explanation for vocal convergence in vampire bats and other echolocating bats is that vocal plasticity evolved for echolocation, and that the calls of echolocating bats might be altered by their auditory inputs as a byproduct of audio-vocal feedback mechanisms that function for echolocation. According to this hypothesis, vocal convergence does not serve a social function, and instead only results from variation in an individual’s exposure to the calls of other individuals.

However, this hypothesis is contradicted by our findings: captive vampire bats that could always hear all colony members in their cages converged with food-sharing partners more than with other cagemates. By contrast, a simple exposure effect would lead only to group-level convergence on calls made by the most vocal bats. Instead, we observed biased vocal convergence towards specific group members depending on the relationships between them. Future work could further assess this hypothesis by comparing how social contact (clustering, allogrooming, food sharing) – as distinguished from acoustic contact alone – influences call similarity.

Vocal convergence might have social causes but not social consequences. An alternative hypothesis is that vocal convergence in vampire bats has social consequences that influence the development of cooperative relationships. For example, it may be a form of behavioural coordination – a key feature of the formation of many social bonds (e.g. coordinated courtship behaviours, vocal exchanges and turn-taking, and reciprocal allogrooming). If partner-specific vocal convergence requires time, it might act as an honest signal of affiliation and investment. On the other hand, if vampire bats can rapidly imitate the calls they hear, then call matching may be used to address specific receivers. For now, the functions of vocal convergence remain unclear.

### Are vampire bats true “vocal learners”?

Definitions of vocal production learning differ among authors. Some authors include modifying extant call types (which is often difficult to distinguish from vocal usage learning), sometimes called “limited vocal learning”. Others emphasize the acquisition of entirely new call types (which is the best possible evidence for vocal learning), sometimes called “complex vocal learning” (26,52–57). Complex vocal production learning is rare among non-human mammals, but several nonhuman species – including cetaceans, pinnipeds, elephants and several species of bats – can copy artificial or novel sounds, including human speech (14,26,52,58–63).

Our findings show that vampire bats are at least ‘limited’ vocal learners, in the sense that social interactions cause increased similarity in either call structure, usage, or both. Although our data are consistent with the hypothesis that adult vampire bats are capable of vocal production learning, further study on variation in call syllables and responses to playbacks would be necessary to assess whether vampire bats are modifying the acoustic features of existing calls in response to auditory input (20,52,53,55). Interestingly, the line between convergence in syllable structure versus syllable usage becomes blurred if contact call variation in vampire bats is not discrete but rather continuously variable. Future work could further delineate contact calls into different syllable types and assess the amount of convergence and individuality within these subjectively defined “call types”.

Evidence consistent with vocal learning has been found in at least 11 species of bats, distributed across eight families (63). Only six of these species have shown *direct* evidence of vocal learning (18,19,59–61,64,65). In the pale spear-nosed bat (*Phyllostomus discolor*) adults can shift the fundamental frequency of their tonal social calls towards a downward-pitch-shifted version of their calls (60), and pups change their isolation calls to resemble their mother’s individual frequency modulation over time (64). In the greater spear-nosed bat (*Phyllostomus hastatus*) individuals changed frequency and temporal characteristics of noisy, broadband “screech calls” to converge on a new group signature as their group composition changed (18). In Taiwanese leaf-nosed bats (*Hipposideros terasensis*), adult bats in the same colony shifted their echolocation calls in the same direction gradually throughout the year, and individuals that joined the colony shifted to match the colony’s mean call frequency (19). In the greater horseshoe bat (*Rhinolophus ferrumequinum*), pups shifted towards the frequency of their mother’s echolocation calls more than to other group members (65). In Egyptian fruit bats (*Rousettus aegyptiacus*), pups raised in normal conditions narrowed their vocal repertoire to fewer adult vocalizations, while experimentally isolated pups lagged dramatically in call development or never developed adult calls (61). In the greater sac-winged bat (*Saccopteryx bilineata*), pups of both sexes learned territorial song syllables from adult males (22,59,66). In general, echolocating bats are remarkably specialized for flexibility in vocal production (52,60) and have evolved sophisticated audio-vocal systems that allow them to adapt their echolocation calls in response to acoustic inputs (67–70). Given the exceptional vocal flexibility of bats, the evidence for vocal learning across the bat phylogeny, and the fact that bat vocal repertoires are large and not well understood (68), it is possible that all bats are capable of vocal convergence or even vocal production learning.

### Conclusion

Manipulations of co-housing in captivity showed that social familiarity causes convergence in contact call structure in vampire bats. Further vocal convergence occurs among familiar female bats that form close social bonds, reflected by dyadic rates of food sharing. Although captive vampire bats in our studies were acoustically exposed to all colony members in their cages, they appeared to converge with food-sharing partners more than other familiar cagemates, suggesting that vocal convergence is not completely explained by mere exposure. The social causes and consequences of this newly discovered vocal convergence have yet to be fully explored.

## Ethics

Procedures with animals were approved by the Institutional Animal Care and Use Committees at the University of Maryland (protocol R-10–63) and the Smithsonian Tropical Research Institute (2015-0915-2018-A9; 2017-0102-2020; 2015-0501-2022), and by the Panamanian Ministry of the Environment (SE/A-76-16 and SEX/A-67-2019).

## Data accessibility

Data and R code for reproducing analyses is available on GitHub (71) and figshare (72): https://github.com/jkvrtilek/call-convergence-2025 10.6084/m9.figshare.29191334

## Declaration of AI use

We have not used AI-assisted technologies in creating this article.

## Authors’ contributions

Julia Vrtilek: data curation, formal analysis, investigation, visualization, writing—original draft, and writing—review and editing; Grace Smith-Vidaurre: data curation, software, formal analysis, investigation, validation, visualization, writing—review and editing; Eric Fosler-Lussier: software, writing—review and editing; Rachel Page: resources, supervision, writing—review and editing; Gerald Carter: conceptualization, data curation, formal analysis, funding acquisition, investigation, methodology, project administration, resources, supervision, visualization, writing—original draft, and writing—review and editing. All authors gave final approval for publication and agreed to be held accountable for the work performed therein.

## Acknowledgements

We thank Gregg Cohen, Imran Razik, and Jerry Wilkinson for help with data collection. We thank May Dixon, Raven Hartman, Virginia Heinen, Tobias Nguyen, Jacob Chisausky, and two anonymous reviewers for feedback that greatly improved the manuscript.

## Supplementary Information (SI)

### Supplementary Methods

#### Variation in contact call structure

We defined contact calls based on a combination of behavioural context and call features. Specifically, we only analysed calls that were produced by a bat that was isolated from group members, had durations of 3-50 ms, and had peak frequencies of at least 10 kHz. Calls were excluded by our analysis pipeline whenever we failed to measure frequencies from the fundamental, indicating low signal-to-noise ratio. We found that removing these calls increased the overall call classification rate for assigning calls to bat.

Within these call parameters, we failed to find any clear evidence for discrete call types despite considerable variation both among and within callers (Figure S1). Given that contact call variation appeared to be graded (continuous) rather than discrete, we decided to take the conservative first step of assessing similarity for all contact calls rather than subjectively defining subtypes and measuring similarity within each type.

The latter approach would also elevate the false discovery rate due to multiple testing and the many possible ways to label types. Future work can assess the degree to which contact call variation is continuous rather than discrete and describe how vocal individuality differs between structurally defined types of syllables.

**Figure S1.**
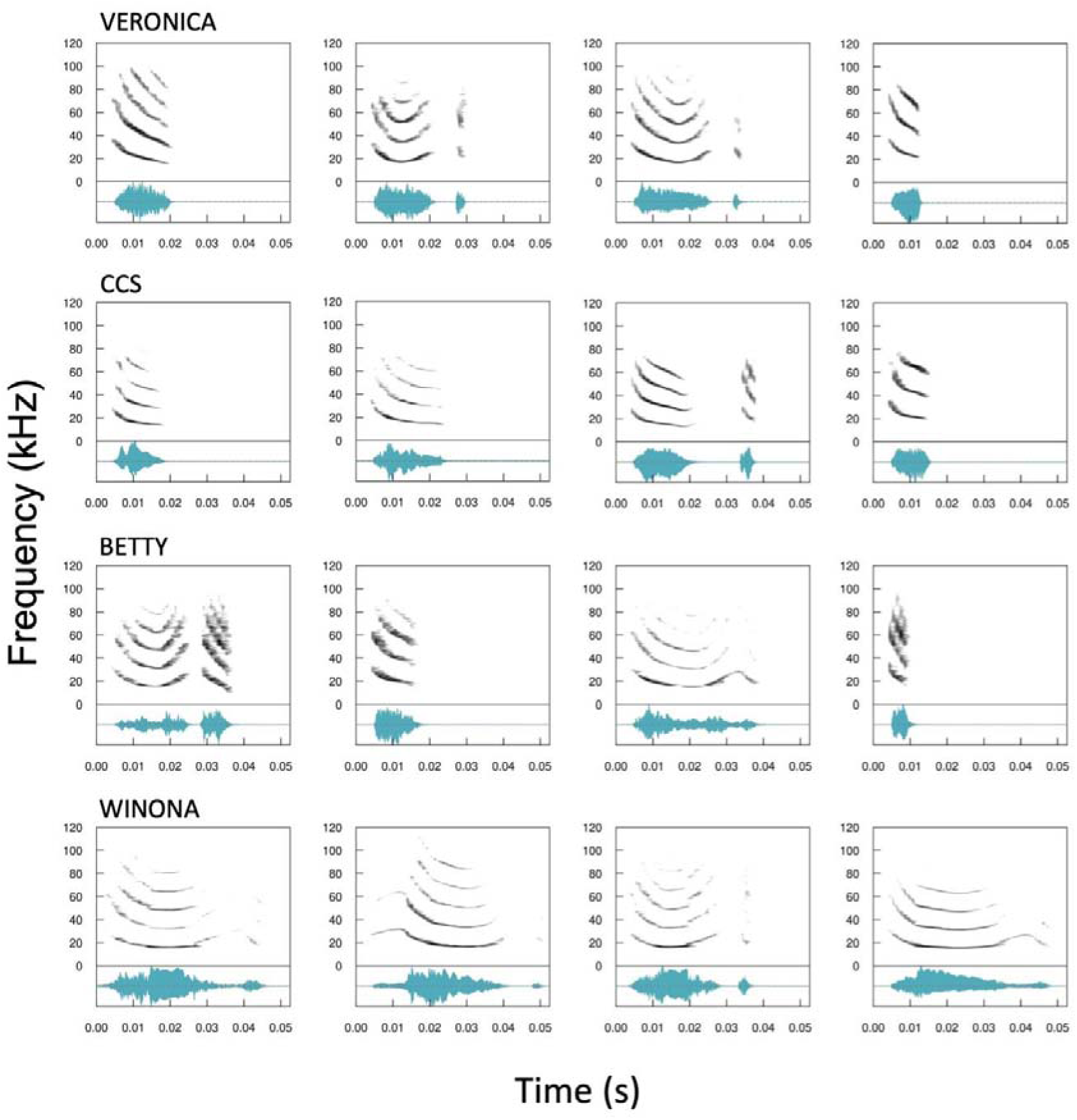
Representative variation in contact calls within and among individual vampire bats. Spectrograms of example contact calls from four female vampire bats (row) across 4 different nights of being individually isolated. Each row represents an individual; the top row is from 2011-2013, the second row is from 2016-2017, and the bottom two rows are from 2019. Recordings in a row are in order from earliest to latest. The variable structure of contact calls suggest that they might convey other information beyond the caller’s individual identity, such as motivational state.

**Figure S2.**
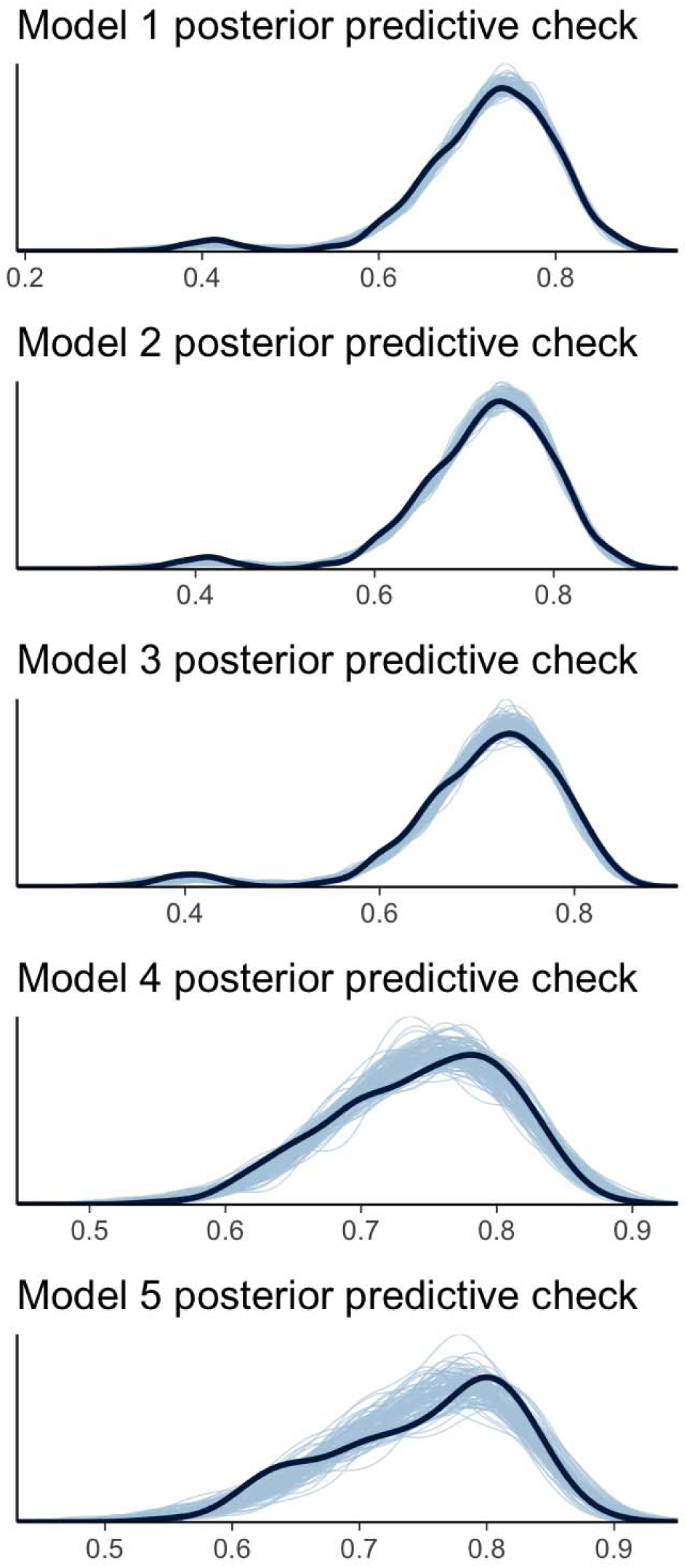
Posterior predictive checks for 5 Bayesian models. Comparison of distribution of observed contact call similarity values (dark black line) to the predicted values from 100 datasets simulated from the fitted model (light blue lines) suggests that the models capture the essential features of the data-generating process.

**Figure S3.**
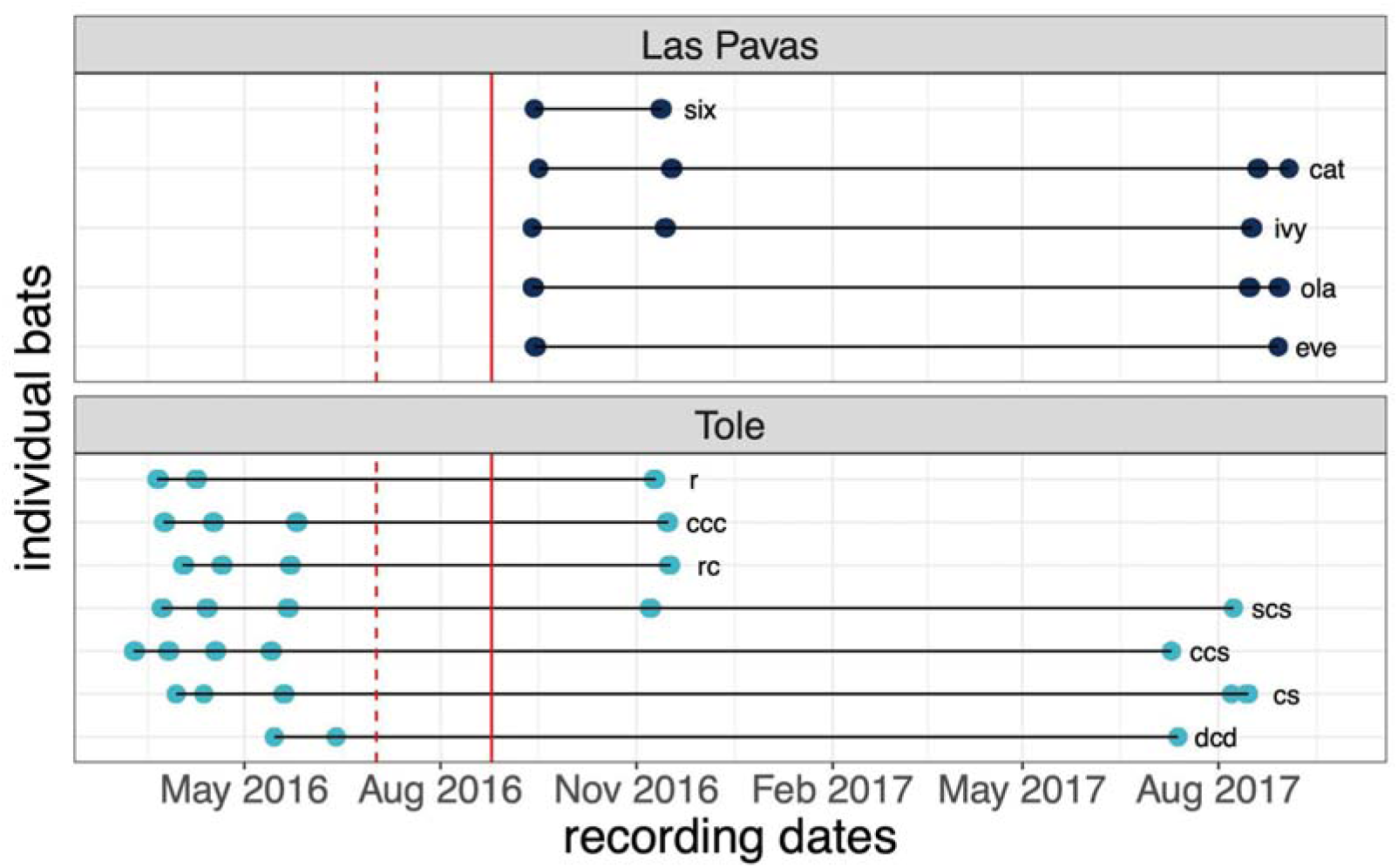
Timeline of call recordings for analysis of call convergence within-pairs over time. Top panel: recording dates (dots) for 5 bats (lines) captured from Las Pavas, Panama. Bottom panel: recording dates for 7 bats from Tolé, Panama. Dotted red line: date of introduction between Tolé and Las Pavas bats in small groups (2-5 bats). Solid red line: date of introduction between all captive bats in large flight cage.

**Table S1.** Bats recorded for this study. Of the 115 bats recorded for this study, 95 produced more than 100 contact calls. The number of bats below adds to 97 because 2 bats from the 2016-2017 colony were recaptured and included in the 2019 colony.

| Years recorded | # bats | Familiarity | Kinship | Origin | Age and Sex |
| --- | --- | --- | --- | --- | --- |
| 2011-2014 | 30 | Long-term captive colony (28,30,31) | Previously estimated using maternal pedigree and microsatellite markers (31) | Multiple zoos | 12 adult F<br>4 subadult F<br>5 adult M<br>9 subadult M |
| 2016 | 4 | Never caged with or introduced to any other bats in the study, but potentially familiar | Unrelated to other bats, but unknown relatedness among each other | Cattle pasture, Chilibre, Panama | 1 adult F<br>3 adult M |
| 2016 | 1 | Never caged with or introduced to any other bat in the study | Unrelated to other bats | Forest, Gamboa, Panama | 1 adult M |
| 2016-2017 | 40 | Co-housed (caged together) for over 10 months (29,73) | Previously estimated using microsatellite markers (73); bats from different sites are unrelated | Cattle pasture, Las Pavas, Panama | 5 adult F |
|  |  |  |  | Captive-born to Las Pavas bats | 1 subadult F<br>1 subadult M |
|  |  |  |  | Tree hollow, Tolé, Panama | 19 adult F<br>7 adult M |
|  |  |  |  | Captive-born to Tolé bats | 6 subadult F<br>1 subadult M |
| 2019 | 22 | Co-housed for 4 months (32) | Bats from different sites are unrelated; kinship unknown for bats from the same site | Same tree, Tolé, Panama | 2 recaptured adult F<br>7 new adult F |
|  |  |  |  | Tree hollow, La Chorrera, Panama | 7 adult F |
|  |  |  |  | Cave, Lake Bayano, Panama | 6 adult F |

**Table S2.** Spectral and temporal measurements used in linear discriminant function analyses. *Duration* through *meanpeakf* descriptions are edited from the *warbleR* (34) and *seewave* (33) documentation. *Maxslope* through *segments* are summary measures of the fundamental from the *soundgen* package (35).

| Measure | Explanation |
| --- | --- |
| duration | Length of signal (in ms) |
| meanfreq | Mean of frequency spectrum (i.e. weighted average of frequency by amplitude within supplied band pass) (in kHz) |
| sd | Standard deviation of frequency (in kHz). |
| freq.median | The frequency at which the signal is divided in two frequency intervals of equal energy (in kHz) |
| freq.Q25 | The frequency at which the signal is divided in two frequency intervals of 25% and 75% energy respectively (in kHz) |
| freq.Q75 | The frequency at which the signal is divided in two frequency intervals of 75% and 25% energy respectively (in kHz) |
| freq.IQR | Frequency range between 'freq.Q25' and 'freq.Q75' (in kHz) |
| time.median | The time at which the signal is divided in two time intervals of equal energy (in s) |
| time.Q25 | The time at which the signal is divided in two time intervals of 25% and 75% energy respectively (in s). |
| time.Q75 | The time at which the signal is divided in two time intervals of 75% and 25% energy respectively (in s). |
| time.IQR | Time range between 'time.Q25' and 'time.Q75' (in s). |
| skew | Asymmetry of the frequency spectrum |
| kurt | Peakedness of the frequency spectrum |
| sp.ent | Energy distribution of the frequency spectrum |
| time.ent | Energy distribution on the time envelope |
| entropy | Product of sp.ent and time.ent |
| sfm | Spectral flatness |
| meandom | Average of dominant frequency measured across the acoustic signal |
| mindom | Minimum of dominant frequency measured across the acoustic signal |
| maxdom | Maximum of dominant frequency measured across the acoustic signal |
| dfrange | Range of dominant frequency measured across the acoustic signal |
| modindx | Modulation index, calculated as the cumulative absolute difference between adjacent measurements of dominant frequencies divided by the dominant frequency range |
| startdom | Dominant frequency measurement at the start of the signal |
| enddom | Dominant frequency measurement at the end of the signal |
| dfslope | Slope of the change in dominant frequency through time (kHz/s) |
| peakf | Frequency with the highest energy from the frequency spectrum |
| meanpeakf | Frequency with highest energy from the mean frequency spectrum (the mean relative amplitude of the frequency distribution of a time wave) |
| maxslope | Maximum slope of fundamental frequency contour |
| minslope | Minimum slope of fundamental frequency contour |
| absminslope | Absolute value of the minimum slope of fundamental frequency contour |
| pos.slopes | Number of positive slopes in the fundamental frequency contour |
| neg.slopes | Number of negative slopes in the fundamental frequency contour |
| turns | Number of turns (changes between positive and negative slopes) in the fundamental frequency contour |
| meanslope | Mean slope of the fundamental frequency contour |
| segments | Number of separate segments in the fundamental frequency contour |

### Supplementary Results

**Figure S4.**
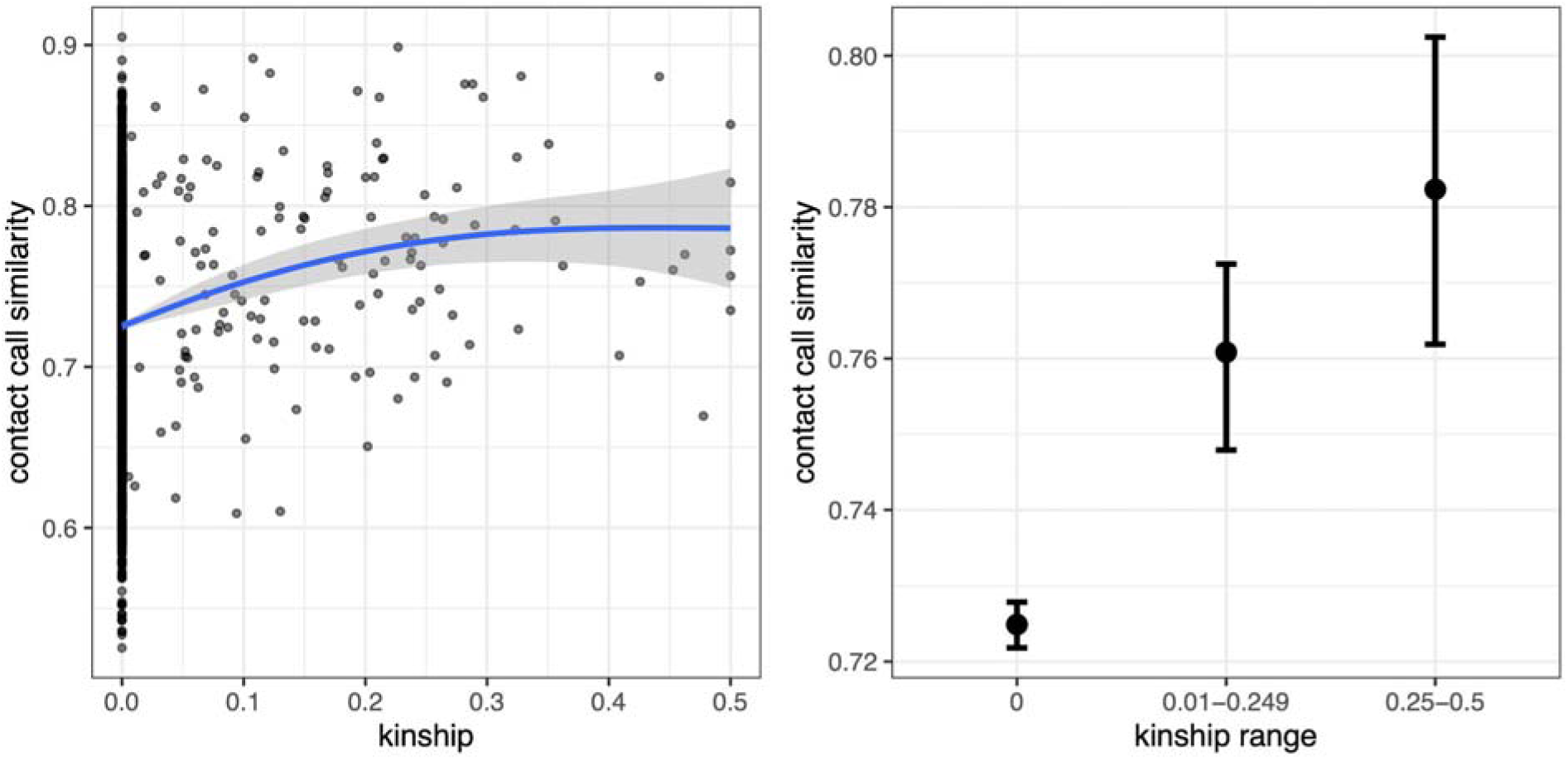
Kinship predicts contact call similarity across pairs with known kinship. Left panel shows how call similarity increases with kinship. Blue line shows a general additive model fit with the ggplot2 R package. Right panel shows mean contact call similarity with bootstrapped 95% confidence intervals (CIs) for three bins of kinship. The 95% CIs treat pairs as independent observations, so these plots are only descriptive; inferences should be drawn from the multi-membership model results (Figure 2b).

**Figure S5.**
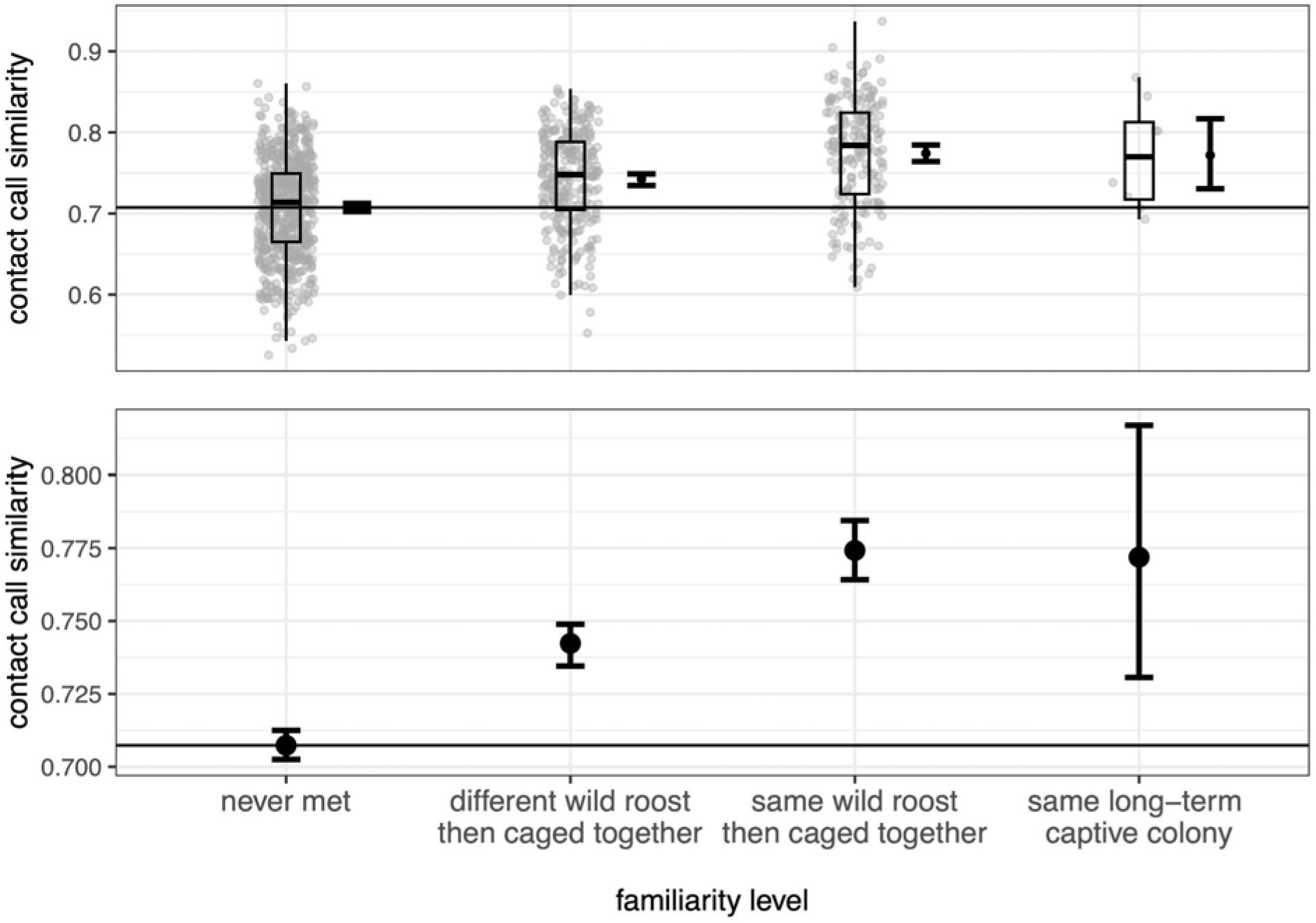
Contact call similarity by levels of familiarity. Top panel shows the distributions and boxplots of the contact call similarity; bottom panel shows means with bootstrapped 95% CIs for contact call similarity. Both are shown at four levels of familiarity (x-axis): (1) unfamiliar nonkin that never met (baseline), (2) previously unfamiliar nonkin captured from different wild roosts then co-housed in captivity, (3) nonkin captured from the same roost then co-housed together in captivity, and (4) familiar nonkin co-housed in the same long-term captive colony. Vocal convergence is demonstrated by the difference from the pairs that never met (horizontal line). The 95% CIs treat pairs as independent observations, so these plots are only descriptive; inferences should be drawn from the multi-membership model results (Figure 2b).

**Figure S6.**
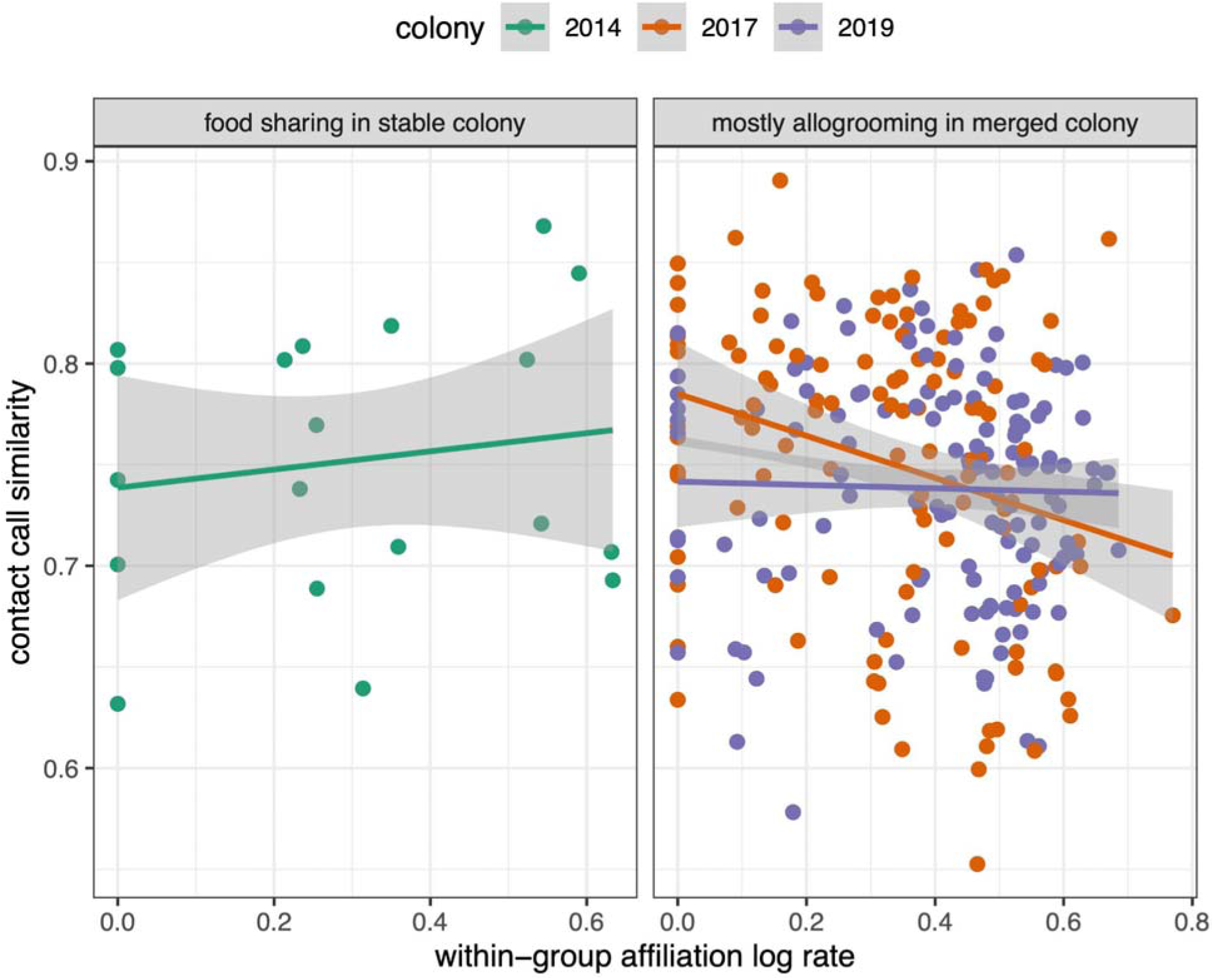
The association between nonkin affiliation rate and nonkin contact call similarity across colonies. Only pairs that were nonkin (r <0.05) are plotted. The 2014 dataset included only food donations to fasted bats in a long-term captive colony (left panel). The 2017 and 2019 colonies were shorter-term colonies that merged samples of bats from different wild populations. The 2017 colony (orange) included mostly allogrooming and some food donations. The 2019 dataset (purple) included only allogrooming in pairs from different sites, because kinship was unknown in pairs from the same site (although including kin pairs does not change the conclusions from these analyses). The 95% CIs on slopes treat pairs as independent observations, so these plots are only descriptive; inferences should be drawn from the multi-membership model results (Figure 2b).

**Figure S7.**
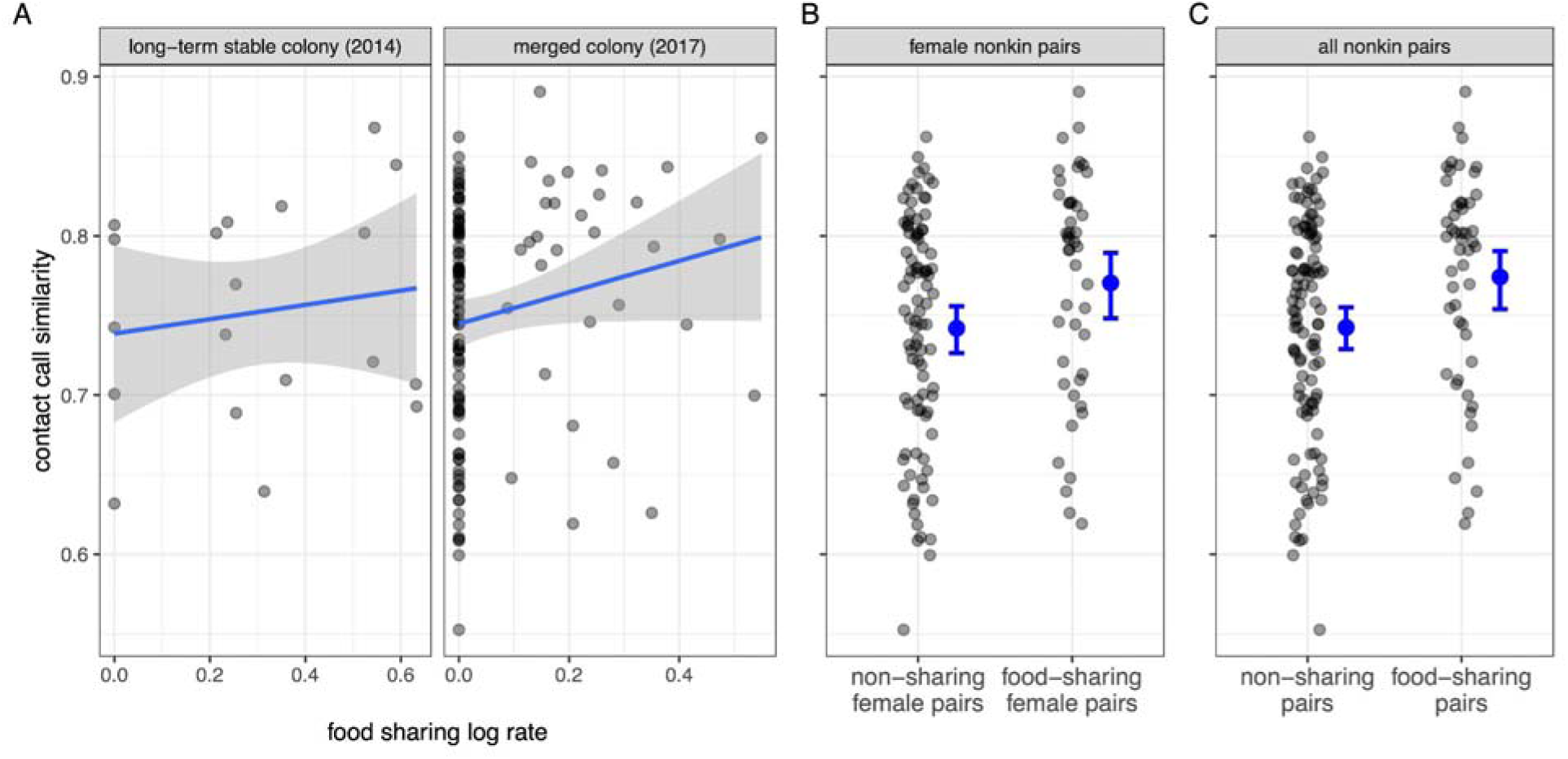
Nonkin pairs that engage in more food sharing tend to have more similar contact calls. Greater food-sharing predicted greater contact call similarity among nonkin females (A). The absence and presence of food sharing is possibly linked to greater contact call similarity in nonkin females (B) and in all nonkin pairs (C). The 95% CIs treat pairs as independent observations so these plots are only descriptive; inferences should be drawn from the multi-membership model results (Figure 2b).

**Table S3.** Model coefficients. Other columns are the estimate of the average (expected) posterior probability, the estimated error (the posterior standard deviation), the 95% credible interval, the R-hat convergence diagnostic (convergence = 1.00), and the Bulk and Tail effect sample sizes.

| Model | Term | Estimate | Est. Error | Lower 95% CI | Upper 95% CI | R-hat | Bulk ESS | Tail ESS |
| --- | --- | --- | --- | --- | --- | --- | --- | --- |
| 1 | kinship (scaled) | 0.037 | 0.005 | 0.028 | 0.047 | 1.000 | 12111 | 11709 |
| 2 | affiliation log rate* (scaled) | 0.025 | 0.006 | 0.013 | 0.037 | 1.001 | 6270 | 9051 |
| 2 | kinship (scaled) | 0.003 | 0.005 | -0.006 | 0.013 | 1.000 | 7686 | 9742 |
| 2 | co-housed (T/F) | 0.148 | 0.013 | 0.123 | 0.173 | 1.001 | 6031 | 8581 |
| 3 | co-housed (T/F) | 0.207 | 0.015 | 0.178 | 0.238 | 1.001 | 7717 | 9579 |
| 4 | affiliation log rate* (scaled) | -0.002 | 0.020 | -0.040 | 0.036 | 1.000 | 8090 | 10802 |
| 5 | food sharing log rate* (scaled) | 0.068 | 0.026 | 0.017 | 0.119 | 1.000 | 10274 | 11916 |

\*Log rate is (natural log (seconds of observation + 1)) / natural log (sampled hours).

